# Recombination landscape dimorphism contributes to sex chromosome evolution in the dioecious plant *Rumex hastatulus*

**DOI:** 10.1101/2021.11.03.466946

**Authors:** Joanna L. Rifkin, Solomiya Hnatovska, Meng Yuan, Bianca M. Sacchi, Baharul I. Choudhury, Yunchen Gong, Pasi Rastas, Spencer C.H. Barrett, Stephen I. Wright

## Abstract

There is growing evidence across diverse taxa for sex differences in the genomic landscape of recombination, but the causes and consequences of these differences remain poorly understood. Strong recombination landscape dimorphism between the sexes could have important implications for the dynamics of sex chromosome evolution and turnover because low recombination in the heterogametic sex can help favour the spread of sexually antagonistic alleles. Here, we present a sex-specific linkage map and revised genome assembly of *Rumex hastatulus*, representing the first characterization of sex differences in recombination landscape in a dioecious plant. We provide evidence for strong sex differences in recombination, with pericentromeric regions of highly suppressed recombination in males that cover over half of the genome. These differences are found on autosomes as well as sex chromosomes, suggesting that pre-existing differences in recombination may have contributed to sex chromosome formation and divergence. Analysis of segregation distortion suggests that haploid selection due to pollen competition occurs disproportionately in regions with low male recombination. Our results are consistent with the hypothesis that sex differences in the recombination landscape contributed to the formation of a large heteromorphic pair of sex chromosomes, and that pollen competition is an important determinant of recombination dimorphism.

## Introduction

The distribution of rates of recombination along chromosomes (recombination landscape [1]) shapes many aspects of evolutionary genetics, including the efficacy of natural selection [2], genome structure [3], and the dynamics of reproductive isolation [4]. Rates of recombination can vary between species, between and within chromosomes, and between male and female meiosis in both dioecious/gonochoristic and hermaphroditic species [5–7]. We refer to this phenomenon as ‘sex differences in the recombination landscape’ [1]. Although sex differences in the rate and distribution of recombination appear to be widespread and variable, the causes and consequences of this variation have only recently been investigated in detail [1,6,8].

Many components of evolutionary processes depend on the sex-averaged rate of recombination. Nevertheless, sex differences in recombination (heterochiasmy) in dioecious populations can have important consequences for the evolution of sex chromosomes. This is because on the sex chromosome restricted to the heterogametic sex (i.e. the Y or W chromosome), sex-specific recombination landscapes entirely control the rate of recombination, and thereby influence the scale of recombination suppression surrounding a sex-determining region (SDR) [1]. Recently, heterochiasmy has been proposed as an important factor in maintaining sexually antagonistic (SA) variants on the sex chromosomes even in the absence of recombination modifiers [9]. In particular, SA alleles can spread more easily through populations because of pre-existing male-specific suppression of recombination, rather than recombination suppression evolving as a secondary consequence of the segregation of SA alleles [9]. Variation among species in patterns of heterochiasmy could thus be an important general determinant of the evolution of sex chromosomes and their turnover [1], potentially contributing to differences among lineages in the likelihood of the formation of heteromorphic sex chromosomes, the maintenance of sexually antagonistic polymorphisms and the size of the SDR.

Several patterns are evident in the characteristics of sexual dimorphism in the recombination landscape. First, although data are limited, many eukaryotes have recombination rates biased towards the tips of chromosomes in male meiosis, whereas female recombination rates are more likely to be either elevated towards the centromeres or are more uniform across the chromosome [1]. In hermaphroditic plants, three of five taxa studied show this pattern [10–12], although in maize there was limited evidence for large-scale differences in recombination between male and female meiosis [13], and in an interspecific cross between *Solanum esculentum and S. pennellii* recombination in male gametes was reduced genome-wide compared with female gametes [14]. Preliminary analysis of genetic maps in the dioecious *Mercurialis annua* do not suggest major sex differences in recombination rates [15], although the genomic context of these maps is still being investigated. In general, however, recombination rate landscape dimorphism has not yet been investigated in dioecious plants, limiting our understanding of its potential role in the evolutionary dynamics of plant sex chromosomes.

Many species have convergently evolved tip-biased recombination in male meiosis [1], but the reasons for this pattern are unclear. Both non-adaptive and adaptive explanations have been proposed. If recombination landscape differences are not adaptive, they may simply result from mechanistic differences between the process of female and male meiosis [1]. Adaptive hypotheses include sexually antagonistic selection favoring tighter linkage between sex-specific genes and regulatory elements [1], selection favoring recombination near the centromere in female meiosis to remove meiotic drive alleles [7], and epistatic haploid selection on male gametes and gametophytes [6]. In plants, evidence that female recombination rates are elevated relative to male recombination rates in outcrossing species compared with selfing species [6] is consistent with models of both female meiotic drive and male gametophytic selection, as both are expected to be more intense with higher rates of outcrossing [6,7]. Disentangling these alternatives is challenging and will require more comparative information on sex-specific recombination in both hermaphroditic and dioecious taxa.

*Rumex hastatulus* is a dioecious, wind-pollinated plant with heteromorphic sex chromosomes [16,17]. Recent genome sequencing combined with high marker-density linkage mapping has revealed that the SDR is embedded within a very large genomic region of highly suppressed recombination [18]. Evidence for similarly large non-recombining regions in the pericentromeric regions of autosomes suggested that pre-existing recombination suppression may have contributed to the formation of large heteromorphic sex chromosomes in *R. hastatulus* [18]. However, this study measured sex-averaged recombination rates, limiting our ability to investigate the potential role of heterochiasmy in sex chromosome formation and maintenance. With earlier evidence for an important role for gametophytic selection on the sex ratio in this species [19,20], the influence of pollen competition in the evolution of the sex chromosomes [21], and indications of frequent male and female transmission distortion in related dioecious *Rumex* taxa [22], there is a strong likelihood that haploid selection in males and/or females may contribute to sex-specific selection favouring sexual dimorphism in recombination landscapes in this system.

Here, we explore the potential importance of heterochiasmy for the evolution of sex chromosomes and test hypotheses concerning the evolutionary forces favouring sex-specific recombination rate differences in *R. hastatulus*. Using a sex-specific linkage map and corrected draft genome assembly, we first determine whether *R. hastatulus* shows evidence for heterochiasmy and other sex differences in recombination landscape and compare this pattern between the sex chromosome and the autosomes. We then examine the correlates of male and female recombination rates genome-wide and use this information to explore the potential role of sexual antagonism, haploid selection, and meiotic drive as evolutionary drivers of sexual dimorphism in the recombination landscape.

## Methods

### Linkage mapping and genome assembly

We generated a mapping population from a cross between a male and female derived from a single population collected in Rosebud, TX [20]. Seeds from the field collection were grown in the glasshouse and at onset of flowering one male and one female individual were randomly paired for a controlled cross to develop the F_1_ generation. Paired plants were immediately moved into miniature crossing chambers [23] to avoid pollen contamination from other plants growing in the same glasshouse, and F_1_ seeds were harvested after maturation. To obtain tissue from F_1_ plants, we sterilized seeds using 5% (V/V) bleach and germinated them on filter paper in refrigerated petri dishes. After germination, we transplanted seedlings into six-inch plastic pots containing a 3:1 ratio of Promix soil and sand and a slow-release fertilizer (Nutricote, 14:13:13, 300mL per 60lbs) and placed them in a glasshouse at the University of Toronto, St. George campus. We watered plants every other day, and their positions on benches were randomized weekly. On day 43 or 44 after transplant, between 10:00 and 12:00, we collected and flash-froze 30mg of leaf tissue for RNA extraction using liquid nitrogen. When plants flowered, we phenotyped for sex. We used Spectrum Plant Total RNA Kits (Sigma Aldrich) for RNA extraction. The sequenced library included 188 individuals: 102 female offspring, 84 male offspring, and three replicate samples of each parent.

For library preparation and sequencing, we sent our RNA samples to the Centre d’expertise et de services Génome Québec (CES, McGill University, Montréal, QC, Canada). CES prepared libraries using NEBNext library prep kits and sequenced them on a NovaSeq6000 S4 PE100. Sequencing resulted in a total of 3.1 billion reads (3,060,570,370), with between 10 and 49 million reads per sample (mean 15,940,471, median 14,548,490). Raw sequence has been deposited on the Sequence Read Archive (SRA) under the accession number PRJNA692236 (embargoed until July 1, 2022 or publication).

We aligned our raw sequencing reads to the *R. hastatulus* Dovetail draft genome assembly [18] using Star 2-pass version 2.7.6 [24,25]. We processed the aligned files to sort, mark PCR duplicates, and (splitNCigar reads) using PicardTools (http://broadinstitute.github.io/picard/) and the Genome Analysis Tool Kit [26].

We initially generated a linkage map using Lep-Map3 [27] from reads aligned to the original *R. hastatulus* draft assembly. The markers could be split into five linkage groups using a LOD score limit of 30 for the initial split followed by a LOD score limit of 34 for the resulting largest linkage group (SeparateChromosomes2). Most of the remaining single markers were put into these groups using a LOD score limit of 25 (JoinSingles2All), totaling about 120,000 markers. We then calculated the marker order for each linkage group with OrderMarkers2 (with default settings).

To improve our linkage map, we reduced redundancy in our genome assembly and constructed a new pseudo-chromosome assembly using the Lep-Anchor [28] software. To obtain reliable linkage maps, we removed the 13 most-recombining individuals from the maps and constructed three independent linkage maps (Lep-MAP3: OrderMarkers2), using only male informative markers (parameter informativeMask=1), only female informative markers (informativeMask=2) and all markers. These maps were used by the Lep-Anchor software.

To reduce redundancy in the genome assembly, we first split the existing Dovetail assembly into contigs based on assembly gaps. Due to a high number of contigs (>44,000), we removed all contigs of < 500bp, full length haplotypes and joined partial haplotypes in windows of five adjacent contigs (link strength was 6 - |distance| - |difference in orientations|, where distance between contig i and j is |i-j| and difference is 0-2 based on how the orientations differ: same=0, one different=1, both different=2). This was done by Lep-Anchor giving it only the alignment chain computed by Haplomerger2 [29] on the WindowMasker [30] soft-masked (contig-split) genome. This allowed us to reduce the number of contigs to about 33,000. All data were lifted to the new contig assembly coordinates using the liftover script and LiftoverHaplotypes module in Lep-Anchor.

We then ran Lep-Anchor (lepanchor_wrapper2.sh) on the final contig assembly using the three linkage maps, new alignment chains (HaploMerger2) and alignments of raw Pacbio sequence aligned by minimap2 [31]. The resulting pseudo-chromosome assembly was used to calculate physical order of linkage map markers and the maps were evaluated (OrderMarkers2 parameter evaluateOrder) in this order to obtain the final linkage maps. After assembly improvement, we compared contig orders between our previous [18] and new maps using custom R scripts incorporating Plotly [32] interactive plotting.

### Recombination rates and transmission distortion

We quantified recombination rates in two ways. First, we described recombination using map lengths in centimorgans (cM) from the maps produced by Lep-Map3. Based on the scale of recombination observed in previous work [18] and the current map, we performed all downstream analyses using 1Mb windows. We also calculated recombination rates as the sum of crossover events per 1Mb window. We first calculated the number of crossovers per site from cM differences using the inverse of the Haldane mapping function [33], then summed crossovers in 1Mb windows. To describe the extent of recombination suppression, we identified the total number of consecutive windows with zero crossovers. We estimated transmission ratio distortion as a likelihood ratio of the deviation from 1:1 transmission of haplotypes from the male and female parent using custom scripts by PR.

### Gene and TE content

We also developed a new annotation using MAKER version 3.01.03 [34]. For our MAKER annotation, we used the soft-masked [30] genome integrated with previously published floral transcriptomes from six individuals [21] and leaf transcriptomes from six populations [17]. Transcripts were assembled with IDBA-tran version 1.2.0 [35] and annotated in four rounds, using the transcripts and the Tartary buckwheat annotation version FtChromosomeV2.IGDBv2 [36] as the evidences for MAKER. We functionally annotated the final annotation based on homology using BLAST version 2.2.28+ [37] and InterProScan 5.52-86.0 [38]. This annotation resulted in 59,121 genes. We also annotated the locations of rDNA repeats using rnammer-1.2 [39]. The parameters used, ‘-S euk’ and ‘-m tsu,ssu,lsu’, indicate that the input reference is a eukaryote, and that we are annotating 5/8s, 16/18s, and 23/28s rDNA.

We produced the TE annotation using the EDTA (Extensive de-novo TE Annotator) version 1.9.7 pipeline [40]. This pipeline combines the best-performing structure- and homology-based TE finding programs (LTR_FINDER_parallel [41], LTR_HARVEST_parallel [42], LTR_retriever [43], TIR-Learner2.5 [44], HelitronScannerv1.1[45], Repeatmodeler-2.0.1 [46] and RepeatMasker-4.1.1 [47] and filters their results to produce a comprehensive and non-redundant TE library [40]. The optional parameters ‘--sensitive 1’ and ‘--anno 1’ were used to identify remaining unidentified TEs with RepeatModeler and to produce an annotation. The ‘EDTA.TEanno.split.gff3’ output file was used as our non-overlapping TE annotation. This file is produced by EDTA by removing overlaps according to the following priorities: structure-based annotation > homology-based annotation, longer TE > shorter TE > nested inner TE > nested outer TE [40].

For all gene content analyses, we used a stringently filtered set of genes to remove gene annotations associated with transposable elements. We first used BEDtools [48] to remove any exons that overlapped a TE, although genes containing both exons that overlapped TEs and exons that did not overlap TEs were retained. We then removed any gene functionally annotated with ‘transpos*’ (transposon, transposase, etc.), ‘ribonuclease H,’ ‘pol poly,’ ‘mitochondri*,’ ‘chloroplast,’ or ‘retrovirus.’ This filtered annotation contained 30,641 genes.

### Differential expression and SNP calling

We performed differential expression analyses using DESeq2 (v. 1.28.1) [49] and our new annotation. For DESeq2 analyses, we aligned reads to the new genome pseudomolecules using STAR version 2.7.6a [49] and generated readcounts using featureCounts (2.02) [50]. Cutoffs for differential expression were as follows: adjusted p-value <0.1, absolute Log2Fold change >1. We identified genes that were differentially expressed between male and female leaf tissues using published leaf RNA sequence data from population samples of the XY cytotype [17], and between male and female floral tissue using published RNA sequence data from the XY cytotype [21]. Genes with fewer than 20 reads across all samples were removed from these analyses. We also identified sequences that were differentially expressed in pollen tissue compared to male leaf tissue, using published sequence data [21]. Finally, we identified sequences differentially expressed in pollen tubes compared to pollen, using pollen from the individuals in the mapping population described [18]. We collected pollen using a kief box (Wacky Willy’s, Victoria, BC, Canada), germinated and grew it in 100μL of media [51] for 24 hours, and flash froze it in LN2. After removing media, we lysed cells and extracted total RNA using Spectrum Plant Total RNA Kits (Sigma Aldrich) for RNA extraction. To identify sex-linked SNPs and fixed differences between the X and Y chromosome (all females homozygous reference or non-reference, all males heterozygous) for our new assembly, we used FreeBayes v0.9.10-3-g47a713e [52] to call SNPs from the population transcriptome data from the XY cytotype (six males and six females) [17] and the crossing transcriptome data from the same cytotype (six male and six female offspring, plus parents) [17]. We filtered the SNPs to exclude any with a SNP quality score of lower than 60, any sites with missing data, and fixed heterozygous SNPs across all samples that likely reflected paralogous mapping.

### Linear modelling predictors of recombination rate

We combined our linkage map data with our annotation, TE annotation, differential expression data and summed and averaged variables in 1-Mb windows to perform analyses of recombination landscape, gene content, and differential expression in R version 4.1.0 [53] in RStudio version 1.4.1717 [54] using the packages dplyr version 1.0.7 [55] and stringr version 1.4 [56]. We performed correlations using R’s built-in cor function, and estimated partial correlations using the package ppcor version 1.1 [57].

To identify factors associated with genome structure that predicted recombination rates and recombination rate differences [58], we created linear models with the following response variables: female crossovers per window, male crossovers per window, sex-averaged crossovers per window, crossover number sex difference per window, and female vs. male biased recombination across window. We fit all responses using generalized linear models with either negative binomial or Tweedie distributions except for female versus male biased recombination, for which we used logistic regression. We performed linear models in glmmtmb version 1.12 [59], evaluated fit using DHARMa version 0.4.3 [60], and compared models using ANOVAs. We performed separate models for each response variable on each chromosome. Scripts are available at https://github.com/joannarifkin/Rumex-sex-specific.

## Results

### Linkage mapping and genome assembly improvement

We identified five linkage groups, consistent with both the karyotype of the XY cytotype of this species [16, 61] and our previous sex-averaged linkage mapping results [18] (table 1). We again identified two apparently metacentric linkage groups (A1 and A2) and three apparently submetacentric linkage groups (A3, A4, XY) based on the patterns of recombination across the chromosomes (figure 1A) and the identities of the scaffolds constituting the linkage groups. We have retained the same autosomal labels across both maps, and they continue to reflect chromosome sizes from largest (A1) to smallest (A4).

**Table 1:** The linkage groups of the revised *Rumex hastatulus* genome assembly, including sex-averaged, male, and female map lengths, length in Mb, gene content, and extent of non-recombining region

|  |  |  |  |  |  |  |
| --- | --- | --- | --- | --- | --- | --- |
| LG | Sex-averaged map length(cM) | Male map length (cM) | Female map length (cM) | Mb* | Number of genes | Largest size of markers with 0 crossover events (Mb), in males/females |
| A1 | 104.57 | 94.624 | 114.516 | 388.3386 (344.5) | 7512 | 238.6 / 94.3 |
| A2 | 91.67 | 79.032 | 104.301 | 278.2766 (260.43) | 7301 | 33.4 / 22.9 |
| A3 | 69.09 | 48.387 | 89.785 | 171.7219 (175.01) | 3923 | 91.5 / 44.3 |
| A4 | 61.29 | 52.688 | 69.892 | 135.2128 (158.24) | 3502 | 62.4 / 30.2 |
| XY | 79.03 | 64.516 | 93.548 | 239.0056 (150.39) | 4752 | 212.1 / 55.3 |

\*Values in brackets indicate the lengths from the previous assembly

**Figure 1.**
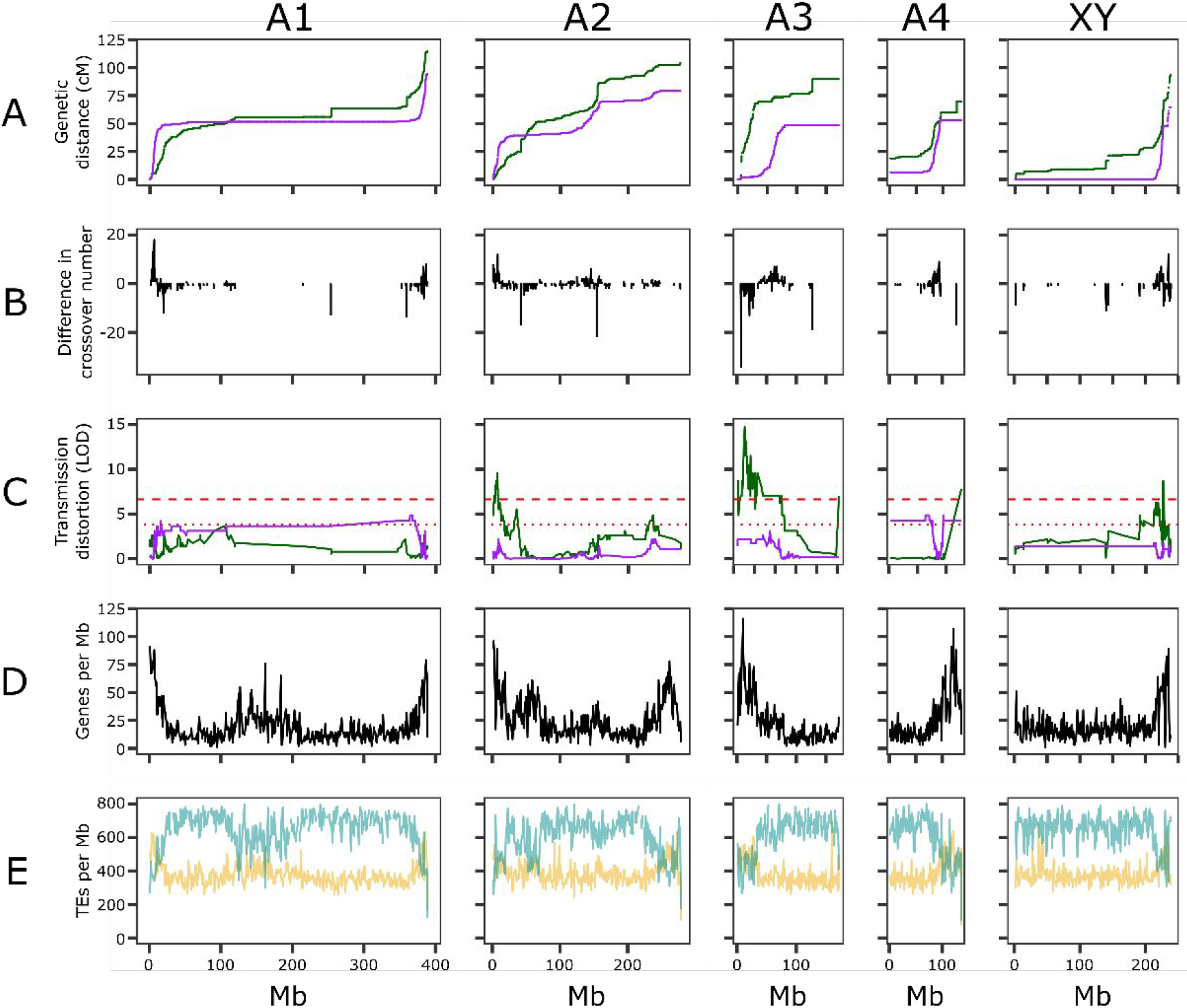
Distribution of recombination, segregation distortion, and gene content in *Rumex hastatulus*. A. Male (purple) and female (green) Marey maps of the chromosomes. B. Difference in crossover number for 1Mb windows along the chromosome (male crossovers per window - female crossovers per window). Positive: male crossover excess. Negative: female crossover excess. C. Segregation distortion for male (purple) and female (green) haplotypes. Dashed red lines indicate significance at 0.05 and 0.01 levels according to a chi-squared test. D. Genes per 1Mb window along the genome. E. TEs per 1Mb window along the genome. Yellow: DNA TEs. Blue: RNA TEs.

Our larger genetic mapping population and improved genome assembly led to considerable improvement in higher-order chromosome-scale scaffolding of the genome of *R. hastatulus*. Our improved genome assembly contained 1.45Gb, a reduction of 0.2Gb from our previous 1.65Gb assembly [18] due to the collapsing of redundant haplotypes (see Methods). For this assembly, 1.212Gb (84%) is now grouped in the five linkage groups (previously 1.08GB, 65% of the previous primary assembly), with the remaining 0.23Gb in smaller contigs. These corrections have substantially increased the size of the assembled sex chromosome, with an additional 88.6 MB of sequence assembled on the sex chromosome, the largest increase of any of the chromosomal scaffolds (table 1). Consistent with this increase, analysis of our past genome assembly showed that only 52% of sex-linked SNPs mapped to the assembled sex chromosome; our new assembly integrated with transcriptomes from independent crossing data [17] now shows that 94% of SNPs showing X-Y segregation patterns map to the sex chromosome.

### Recombination rates

As in our previous study, we found that recombination was unevenly distributed across the genome, with very large non-recombining regions on all chromosomes (figure 1). We identified clear evidence of heterochiasmy (table 1, figure 1A). Male map lengths were shorter than female map lengths: across chromosomes, female map length was 1.4x male map length (table 1, figure 1A), and the sex chromosome was not an obvious outlier for this metric. Males also had longer blocks of non-recombining windows across all chromosomes. To summarize this pattern, we identified the longest stretches of markers on each chromosome with zero crossovers. On the autosomes, males had runs of non-recombining windows approximately twice as long as those of females, with male-specific non-recombining regions as large as 238MB (table 1). By this measure, the sex chromosomes were an exception: the largest run of male-specific non-recombining windows on the sex chromosome was four times the length of the longest run of female-specific non-recombining sequence (table 1). Thus, although all chromosomes exhibited reduced male recombination rates, the XY chromosome showed the largest region of differentially suppressed recombination between the sexes despite not being the largest chromosome. Overall, male and female recombination rates in *R. hastatulus* were only weakly correlated (*r*=0.333, correlation of male and female crossover number across 1Mb windows across all chromosomes; figure 2, table S1).

**Figure 2:**
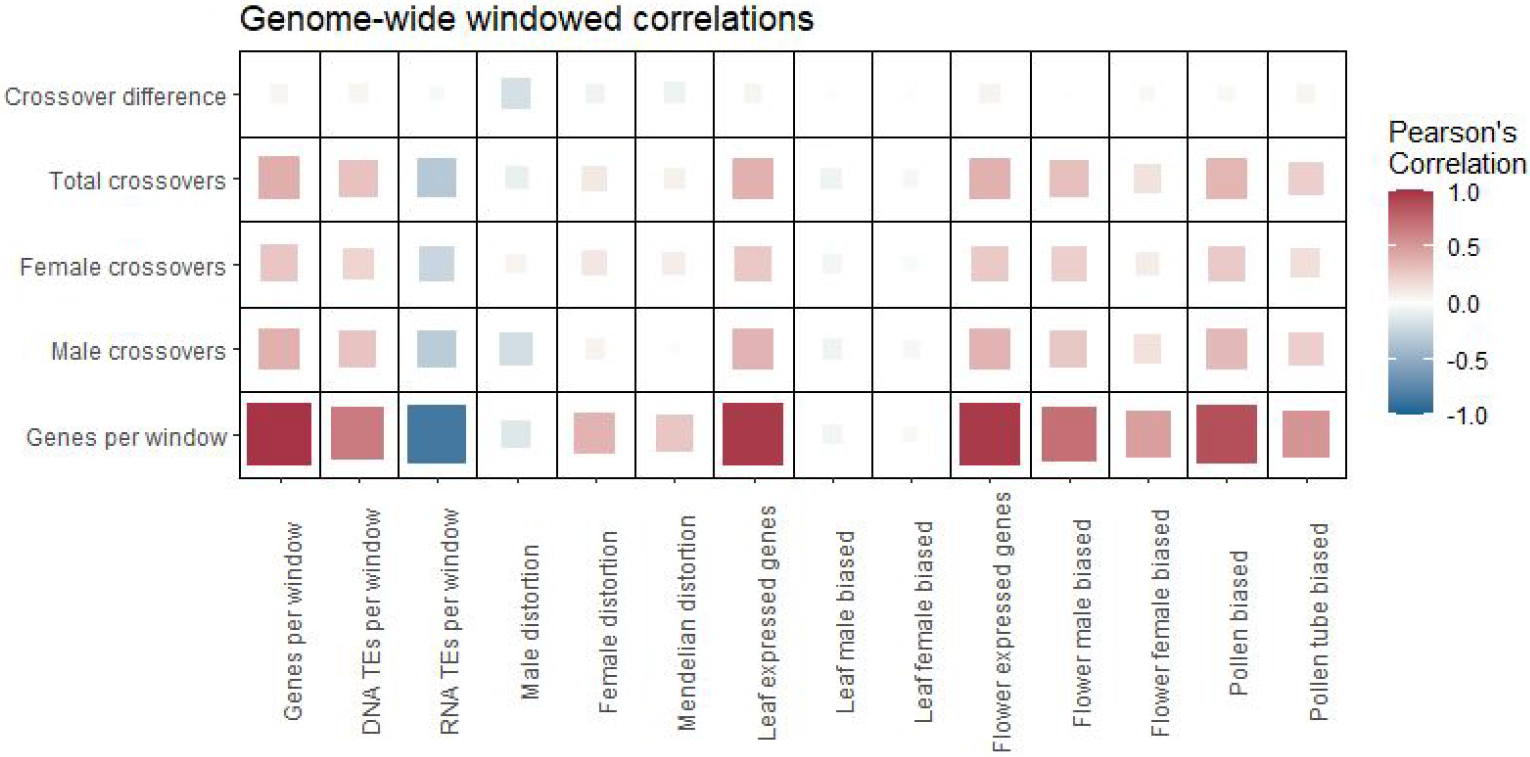
Correlations among genome window characteristics across the whole genome of *Rumex hastatulus*. Colours and sizes correspond to the strength and direction of the correlations.

The extent of sex differences in recombination varied both along and between chromosomes (figure 1B). Chromosomes A1, A2, and the sex chromosome conformed to the common pattern of more tip-focused recombination in males, but the submetacentric chromosome A3 showed female-biased recombination in the more highly recombining end, and A4 appeared to have low-recombination regions at both ends. This differs from the previous linkage map, likely because of the difficulty of positioning low-recombination regions. In general, pericentromeric regions showed reduced recombination in both sexes, but female map lengths were larger in these regions and showed apparent hotspots of recombination with large jumps in centimorgan position (figure 1). All five chromosomes had regions of female-biased and male-biased recombination, as well as shared recombining and non-recombining regions (figure 1B). This complex pattern creates a highly heterogeneous recombination landscape.

### Gene and TE content

As in our previous study, we found that genes were generally concentrated in high-recombination regions (figure 1D). However, A1 contained one gene-dense yet low-recombination region (∼100Mb-200Mb). RNA (class 1) TEs were concentrated in low-recombination regions, whereas DNA (class 2) TEs were concentrated in high-recombination regions (figure 1E). Despite the reduced gene density in low-recombination regions, the extent of high recombination-suppressed regions means that a large fraction of genes in the genome (approximately 37%) are in these large regions with no male recombination. Ribosomal genes were concentrated mostly on A3 (62 rDNA features annotated) and A4 (28 rDNA features annotated). rDNA genes occurred in the first 50Mb of A3 (5S subunit sequence) with additional rDNA sequence located around 130Mb of A3 (18S and 28S subunit sequence) and in the first 3 Mb of A4 (18S and 28S subunit sequence). These rDNA locations are consistent with past cytological findings [61], further confirming our identification of the A3 and A4 chromosomes.

### Characterization of the sex-determining (SDR) and pseudo-autosomal regions

With female and male recombination separated, it is clear that the male-specific non-recombining region on the sex chromosome in *R. hastatulus*, i.e. the SDR, is extensive. In particular, our linkage mapping suggests that the SDR is as large as 209MB (14% of the total assembly: table 1, figure 1), including as many as 3595 genes (12% of the total filtered annotated genes). The gene-dense recombining pseudoautosomal region of the sex chromosome is similarly the smallest male recombining segment of any chromosome, representing only approximately 13% of the physical size of the chromosome. Note that the ‘true’ SDR may be narrower and the pseudoautosomal region larger, since rare male recombination may have gone unobserved in our cross. However, BLAST [37] searches of our sex-linked transcripts that have at least one fixed difference between the X and Y chromosomes from a population sample [17] are found across most of the length of this nonrecombining region (from 1.6 MB to 208.5 MB), and fixed differences mapped onto the chromosome are common across the first 210 MB (figure S1) suggesting that most of this region is nonrecombining and linked to the SDR.

### Transmission ratio distortion

We identified loci with biased haplotype transmission through either paternal or maternal inheritance based on the haplotype reconstructions from Lep-Map 3. Transmission ratio distortion varied between chromosomes (figure 1C, figure S2, table S2). More sites experienced biased transmission through maternal (276) than paternal (48) inheritance, across a larger fraction of the genome. On A1, 18 sites were significantly distorted in transmission from males using a 0.05 cutoff in a chi-squared distribution (LOD > 3.841) and on A4, 30 sites were significantly distorted in transmission from males with a 0.05 cutoff. Although deviations from 1:1 male haplotype transmission occurred on other chromosomes, there were no significant male-distorted sites on A2, A3, or the sex chromosome. In contrast, female haplotype distortion was extensive on A2 (91 sites in females at 0.05, 38 at 0.01 cutoff of LOD > 6.635), A3 (108 sites at .05 cutoff, 88 at 0.01), and the sex chromosome (76 sites at a 0.05 cutoff, 26 at 0.01) but negligible on A1 (0 sites) and A4 (1 site at 0.01).

### Correlates of recombination rate differences

Across the genome, the number of genes, leaf- and flower-expressed genes, and DNA (Class 2) TE density were all positively correlated with recombination rate, and RNA (Class 1) TE density was negatively correlated with recombination rate (figure 2, table S1, figure S3-S6). Male crossover number and female crossover number were both positively correlated with gene density, but this correlation was stronger for male crossovers (male Pearson’s *r*=0.390, female Pearson’s *r*=0.278). However, the correlations between transmission ratio distortion and crossover number were in opposite directions: in females, more distorted regions were also more recombining (*r*=0.111) whereas in males distortion was negatively correlated with crossover number (*r*=-0.197), reflecting the fact that male transmission distortion signals were enriched in the large non-recombining regions of male meiosis (figure 1). Correlations varied in strength between chromosomes, with notable differences in correlates of transmission ratio distortion and recombination rate difference, both of which varied in magnitude, position, and direction along and between chromosomes (figure 2, figure S7, table S3). The difference in crossover number between the sexes was most strongly correlated with signals of male distortion (figure 2), where regions of particularly low male crossover number represented regions with larger signals of male distortion. We also estimated partial correlation coefficients controlling for gene density, which was consistently correlated with many genomic variables (figure S8, figure S9, tables S4-S5). Correlations between distortion and crossover number in both males and females persisted after controlling for gene density, and varied in direction across chromosomes (figure S9). However, all chromosomes except A2 showed a consistent negative correlation between male-biased recombination and male transmission distortion, even when gene density was controlled.

### Linear models of recombination rate differences

We used generalized linear models to identify predictors of sex-specific and sex-averaged recombination rates, whether recombination was male- or female-biased, and the magnitude of the recombination rate difference in *R. hastatulus*. Full modeling results are available in table S6 and table S7.

In our linear models, the variables that significantly predicted male recombination rate varied between chromosomes (table 2A). Number of genes predicted increased male recombination rate on three chromosomes (as well as a fourth in some models), and position along the genome emerged as an important predictor on two submetacentric chromosomes, suggesting that distance from centromere predicts male recombination (figure 1). RNA TE count also appeared to play a role, but in inconsistent directions, predicting increased male recombination on two chromosomes and decreased male recombination on a third; this pattern also appears in the partial correlations with gene density removed (figure S9). Transmission ratio distortion predicted reduced male recombination rate on three chromosomes (A2, A3, and A4), and predicted increased male recombination on A1. Either pollen bias or pollen tube bias predicted male recombination rate on three chromosomes (negatively on A4 and the sex chromosome, positively on A3).

**Table 2.**
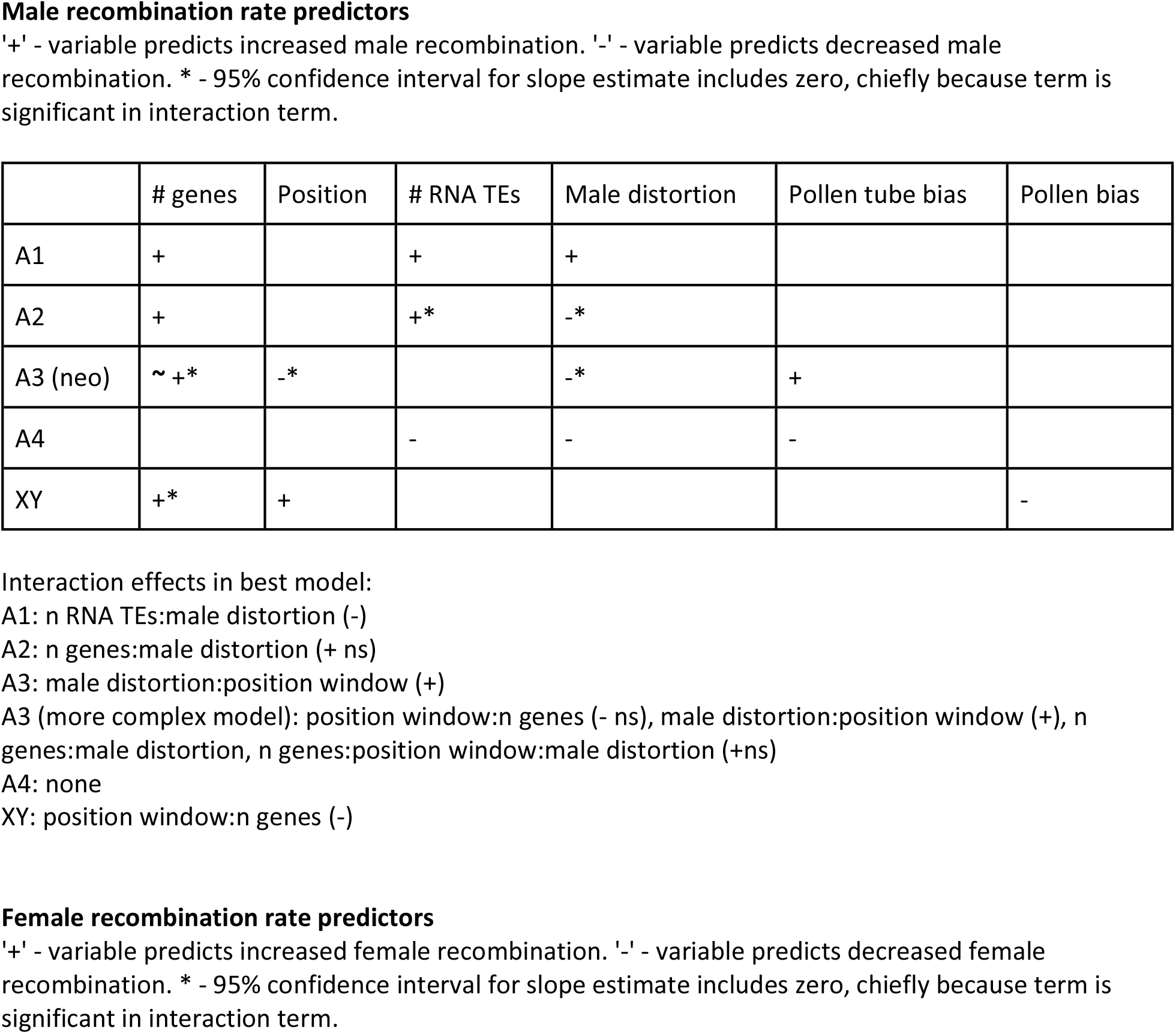
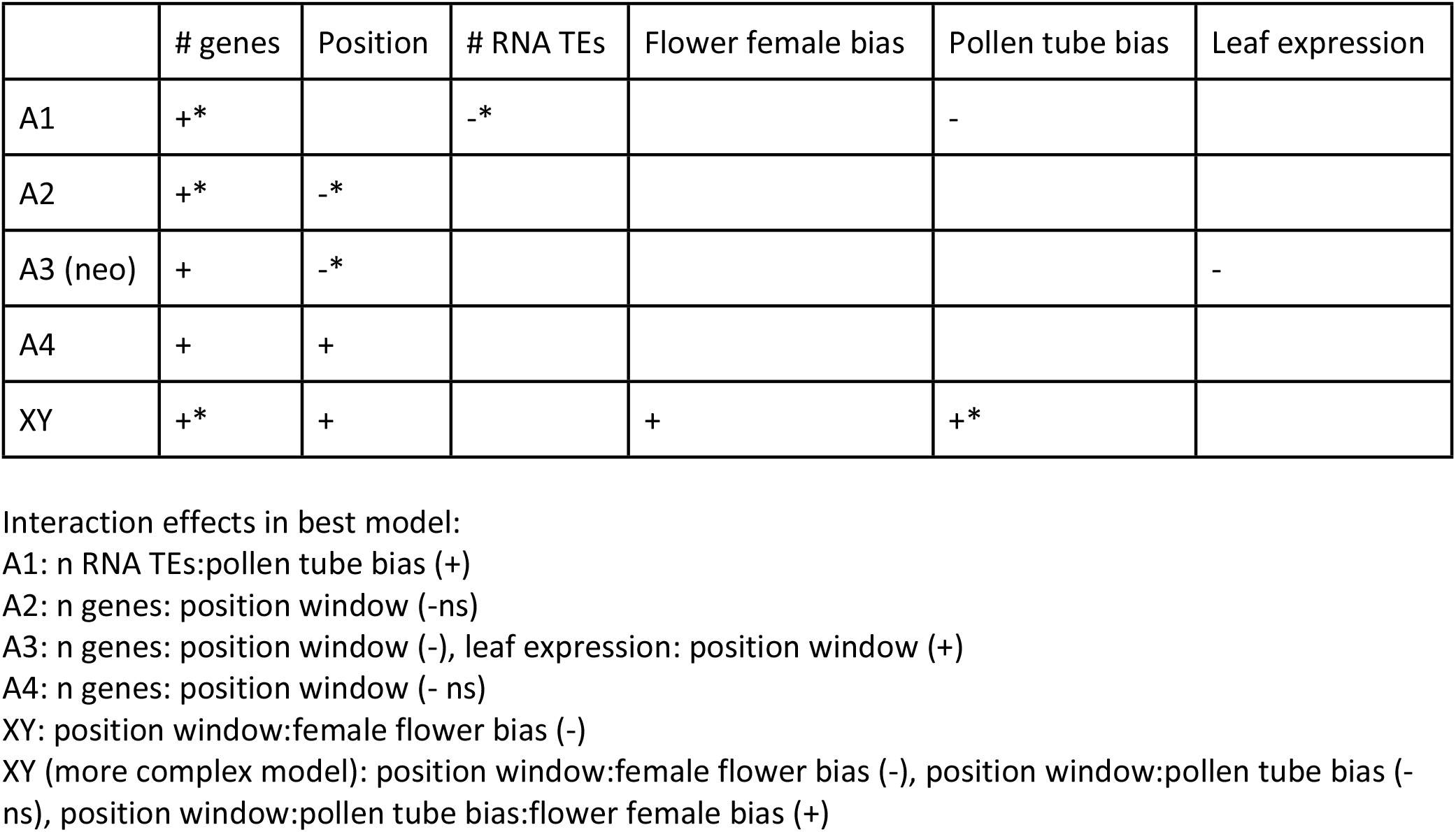
Variables identified as significant predictors of recombination in linear models of windows of the *Rumex hastatulus* genome.

In contrast, the predictors of female recombination are more consistent across chromosomes (table 2B). Number of genes per window positively predicted female recombination rate on all five chromosomes, and position relative to centromere affected female recombination rate on four chromosomes. Only one other variable, pollen tube-biased expression, predicted female recombination rate on more than one other chromosome. Female transmission ratio distortion did not emerge as a significant predictor of recombination rate on any chromosome.

Predictors of both sex-averaged recombination and of the magnitude of the recombination rate difference between males and females reflected predictors of male and female recombination rate independently. Number of genes, position along the chromosome, number of RNA TEs, female-biased floral expression, pollen tube-biased expression all appeared as significant predictors, but their importance and direction varied between chromosomes (table S6A, S6B).

Finally, we used logistic regression to identify variables that predicted whether windows exhibited female or male recombination bias (table S6C). Our logistic regressions also suggested considerable variation in the factors that predicted sex differences in recombination. Number of genes per window was an important predictor for four out of five chromosomes, but in variable directions, predicting both male bias (A1, XY) and female bias (A2, A3) in recombination.

## Discussion

The main findings of this study are consistent with the general pattern of extensive recombination suppression that we previously inferred based on sex-averaged recombination in *R. hastatulus* [18]. However, our sex-specific maps show that recombination suppression is not evenly distributed between males and females, and that the observed very large pericentromeric regions of suppressed recombination are particularly influenced by highly suppressed male recombination. Across all chromosomes, females recombine more frequently than males and males have much larger non-recombining blocks than females. Our results are in line with several studies of hermaphroditic plants, as well as other eukaryotes [1], with the very large male-specific non-recombining regions that we report making this an extreme case.

However, *R. hastatulus* does not strictly follow the common eukaryotic pattern of tip-focused male recombination [1] across all chromosomes. The recombination landscapes of both the metacentric and submetacentric chromosomes suggest greater variation in the distribution of recombination than simply less male recombination in the centres and more at the chromosome ends, with highly recombining regions scattered along chromosomes (particularly A2, A3, and A4). Thus, male recombination is more concentrated, but not always at the tips of the chromosome. These differences among chromosomes may reflect an ongoing history of chromosomal rearrangements in the genus and additional patterns of chromosome structure such as the locations of centromeres and rDNA clusters [61].

The pattern of larger non-recombining regions in males is consistent with an evolutionary bias toward the evolution of male heterogamety and XY sex chromosomes in *Rumex*. In particular, suppressed recombination can facilitate the maintenance and invasion of sexually antagonistic variants linked to sex-determining regions [62], and sex-determining regions that evolve in large pericentromeric non-recombining regions, especially sex-specific ones, may contribute to the evolution of sex chromosomes [63]. The existence of male-specific non-recombining regions may thus facilitate the evolution of XY rather than ZW sex chromosomes [1]. Given our observation of no crossovers in male meiosis over very large fractions of each chromosome, it is possible that recombination suppression was ancestral to the evolution of dioecy in the genus, and that subsequent recombination modifiers did not evolve following the origin of the SDR. The observed size of the SDR, which is over 200 MB using population-validated sex-linked genes and includes over 14% of the assembled genome and over 3500 genes, is much larger than those recently reviewed to date in plants [64], although larger population samples of *R. hastatulus* should be used to test for very rare recombination between the X and Y, and the sex-linked region of *S. latifolia* may be even larger [65]. Although we cannot rule out a role for subsequent recombination modifiers, particularly because the sex chromosomes show the most extreme size dimorphism of the nonrecombining region (table 1), our results do suggest that sex differences in heterochiasmy may have played an important role in determining the large size of the SDR facilitating the evolution of large heteromorphic sex chromosomes in this system. Comparative studies of heterochiasmy in both hermaphroditic and other dioecious species in *Rumex* will be important to further assess the extent to which ancestral heterochiasmy and derived changes in recombination rates have contributed to sex chromosome evolution in this lineage.

Models for the evolution of heterochiasmy due to male haploid selection and female meiotic drive both predict lower overall male recombination rates and higher female recombination near centromeres [1,6,7], as we observed in *R. hastatulus*. Our mapping population provided evidence for both male and female transmission ratio distortion, which varied within and among chromosomes and between the sexes. Overall, more sites displayed significant distortion in female than in male transmission, but significant regions of both types of distortion were observed across the genome. Transmission ratio distortion through female inheritance is generally thought to be consistent with female meiotic drive. In contrast, transmission ratio distortion through male inheritance may result from haploid (pollen) competition [66].

Nevertheless, zygotic distortion (i.e., differential seed germination or seedling survival) could also lead to transmission ratio distortion, and may result from alleles inherited from either parent [66]. In our study, we genotyped reproductive adults rather than pollen or seeds, which conflates several opportunities for natural and sexual selection causing biased transmission. Zygotic selection may be particularly likely to explain our observed female distortion, since these regions were not focused on low-recombination centromeric regions, where meiotic drive is expected to act [7]. In contrast, signals of male transmission distortion are particularly enriched in regions of low male recombination and high sex bias in recombination (figure 1 and figure 2, table 2), suggesting that haploid selection in males may be an important selective pressure for sex differences in recombination. Distorted regions can vary widely between populations of the same species [67], so distortion in a single cross should be interpreted with some caution. Furthermore, patterns of biased pollen or pollen tube expression do not show similarly consistent enrichment in regions of low male recombination except for the sex chromosome (table 2, figure S6), although direct observation of transmission distortion likely provides a stronger indicator of loci involved in pollen competition. Similarly, although we did not identify evidence consistent with ongoing recombination increases to counter meiotic drive, the overall pattern of increased and more centromere-biased recombination is still consistent with a history of selection eliminating meiotic drive alleles. With those caveats, our results do provide some evidence in accord with the hypothesis that pollen competition may play an important role in sex differences in recombination.

We used both pairwise correlations and regression models to investigate various genomic correlates of male and female recombination and sex bias in recombination, in order to further explore other possible factors favouring sex differences in recombination. On a genome-wide scale, both male and female recombination rates in *R. hastatulus* are consistent with widely observed patterns that genes and Class 2 DNA TEs concentrate in high-recombination regions and Class 1 RNA TEs concentrate in low-recombination regions (figure 2; [3], [56]). These patterns are consistent with recombination rates in plants occurring preferentially upstream of genes, and with epigenetic silencing of retrotransposable elements causing a reduction of recombination, although they could also be explained if transposable elements preferentially accumulate in regions of low sex-averaged recombination [3].

Because the recombination landscape varied widely between the chromosomes of *R. hastatulus*, we also identified chromosome-specific predictors of recombination rates and recombination rate bias using linear models and partial correlations controlling for gene density. At this more granular scale, a different and more complex picture emerges (table 2, figure S8, S9). In particular, our logistic regression models of recombination bias direction support the possibility that different mechanisms may be contributing to variation in recombination rates on different chromosomes. Both number of genes per window and position along the genome predicted female-biased recombination on some chromosomes and male-biased recombination on others. Aside from physical position and gene density, different diverse factors predicted both male and female bias on different chromosomes, including RNA TEs, number of genes expressed in different tissues, sex-biased floral expression, pollen-biased expression, and transmission ratio distortion (table 2C). This finding suggests a general picture in which gene density and proximity to centromere shape recombination on a ‘global scale’, but variation in gene content and haploid selection may lead to region-specific selective forces acting on both male and female recombination rates. Consistent with this, a recent comparative study in fish [68] found that sex differences in recombination are labile at the species level, but do not present clear trends consistent with adaptive hypotheses across species.

## Conclusions

Our study has provided some of the first evidence of sex differences in recombination and identified one of the largest known SDRs in a dioecious plant species, or in fact in any eukaryote [65]. We identified both genome-wide and chromosome-specific factors predicting sex differences in recombination, and also found evidence consistent with a role for male gametophytic selection in driving these differences. Future work in this system will allow more precise dissection of the genetic and evolutionary mechanisms favouring sex differences in recombination landscapes. In particular, exploring both sex-specific eQTL positions and further study of transmission ratios in pollen and seeds will allow us to further differentiate between sexually antagonistic cis epistasis in diploids and epistasis in haploids [1]. Finally, characterizing sex-specific recombination landscapes of hermaphroditic *Rumex* species should make it possible to determine whether these sex differences in recombination landscape did indeed precede and promote the evolution of a heterogametic XY sex determining system with a very large SDR, as we have hypothesized.

## Supporting information

A3 windowed correlations

A3 windowed partial correlations

A4 windowed correlations

A4 windowed partial correlations

Table_S1_Genome_wide_windowed_correlations

Table_S2_Genotypes_for_all_markers

Table_S3_all_chromosomes_windowed_correlations

Table_S4_Genome_wide_windowed_partial_correlations

Table_S5_all_chromosomes_windowed_partial_correlations

Table_S7_linear_model results

XY_windowed_correlations

XY_windowed_partial_correlations

A1_windowed_correlations

A1_windowed_partial_correlations

A2_windowed_correlations

A2_windowed_partial_correlations

## Acknowledgments

We thank University of Toronto undergraduate students Victoria Marshall, Claire Ellis, Deanna Kim, and Madeline Jarvis-Cross for technical assistance, Bill Cole and Tom Gludovacz for glasshouse support, and Brechann McGoey for crossing chamber development. This research was supported by Discovery grants from the Natural Sciences and Engineering Research Council of Canada to SCHB and SIW. JLR was supported by an EEB post-doctoral fellowship.

## Data accessibility statement

Raw sequence has been deposited on the Sequence Read Archive (SRA) under the accession number PRJNA692236 (embargoed until July 1, 2022 or publication). Our new genome assembly, transcriptome annotation, rDNA annotation, and TE annotation have been deposited in the CoGe comparative genomics platform at https://genomevolution.org/coge/GenomeInfo.pl?gid=62326. Scripts used in the analyses have been deposited on Github at https://github.com/joannarifkin/Rumex-sex-specific.

## Notes

***Data*** *It is a condition of publication that data, code and materials supporting your paper are made publicly available. Does your paper present new data?:* Yes

***Conflict of interest*** I/We declare we have no competing interests

https://github.com/joannarifkin/Rumex-sex-specific

### Competing Interest Statement

The authors have declared no competing interest.

