## Supplement for "Recombination landscape dimorphism contributes to sex chromosome evolution in the dioecious plant *Rumex hastatulus*"

### Linear modeling results

#### Table S1

Correlations between selected genomic features in 1Mb windows, genome-wide. (Attached.)

#### Table S2

Genotypes at all markers. First position indicates allele inherited from father, second position indicates allele inherited from mother. (Attached)

#### Table S3

Correlations between selected genomic features in 1Mb windows, by chromosome. A: A1. B: A2. C: A3. D: A4. E: XY. (Attached.)

#### Table S4

Partial correlations between selected genomic features in 1Mb windows, genome-wide, controlling for gene density. (Attached.)

#### Table S5

Partial correlations between selected genomic features in 1Mb windows, controlling for gene density, by chromosome. A: A1. B: A2. C: A3. D: A4. E: XY. (Attached.)

#### Table S6

Summary of linear modeling results for total recombination rate, magnitude of recombination rate, and direction of recombination rate bias.

##### A. Total recombination rate predictors

'+' - variable predicts increased number of crossovers per window. '-' - variable predicts increased number of crossovers per window. * - 95% confidence interval for slope estimate includes zero, chiefly because term is significant in interaction term.

|  | # genes | Position | Pollen tube bias | Pollen bias | Flower female bias | Male distortion | Female distortion |
| --- | --- | --- | --- | --- | --- | --- | --- |
| A1 | +* |  | + |  |  | - | - |
| A2 | + | +* |  | + |  |  |  |
| A3 (neo) |  | -* |  |  |  | + | - |
| A4 | + | + |  |  |  | - |  |
| XY (a)  XY (b)^1^ | +*  + | -  - |  |  | + |  | - |

Interaction effects in best model:

A1: n genes:male distortion (+)

A2: position window:n genes (-), pollen bias:n genes (-)

A3: position window:male distortion (-), position window:female distortion (+)

A4: position window:n genes (-)
XY (a): position window:female distortion (+)

XY (b): position window:flower female bias (-)

^1^On both A4 and the sex chromosome, both a model including distortion that only fitted recombination in distorted regions (model a) and a model including flower female bias that fitted recombination chromosome-wide (model b) were effective, and could not be directly compared because they included different datasets. On A4, the chromosome-wide model exhibited clear evidence of problems with fit (Kolmogorov-Smirnov and residual quantile deviation tests <0.01), so we selected the model including distortion. On the sex chromosome, however, both models fit their respective datasets well.

##### B. Recombination rate difference predictors

'+' - variable predicts increased difference between male and female recombination. '-' - variable predicts increased difference between male and female recombination. * - 95% confidence interval for slope estimate includes zero, chiefly because term is significant in interaction term.

|  | # genes | Position | # RNA TEs | Flower female bias | Pollen tube bias | Male distortion |
| --- | --- | --- | --- | --- | --- | --- |
| A1 | -* |  | - |  | - |  |
| A2 | -* | - | -* |  |  |  |
| A3 (neo) |  | +* |  | - |  | + |
| A4 | + | + |  | -* |  |  |
| XY | + | + |  | + |  |  |

Interaction effects in best model:

A1: n genes:RNA TEs (+), n genes:pollen tube bias (+), RNA TEs:pollen tube bias (+)

A2: n genes:RNA TEs (+)

A3: position window : male distortion (-)

A4: position window:n genes (-), position window:flower female bias (+)

XY: position window:flower female bias (-)

Another model with female distortion was better than other models on the distortion subset of the data, but exhibited convergence problems because of the small dataset

##### C. Binary recombination rate difference predictors

'm' - variable predicts greater male bias in recombination. '-' - variable predicts greater male bias in recombination. * - 95% confidence interval for slope estimate includes zero, chiefly because term is significant in interaction term.

|  | # genes | Position | # DNA TEs | # RNA TEs | Leaf expressed genes | Flower expressed genes | Male-biased flower genes | Female-biased flower genes | Pollen bias | Pollen tube bias | Male distortion | Female distortion |
| --- | --- | --- | --- | --- | --- | --- | --- | --- | --- | --- | --- | --- |
| A1 | m* |  |  | f* |  |  |  | f |  |  |  | f* |
| A2 | f* |  |  |  |  |  | f |  |  |  | m* |  |
| A3 (neo) | f* | f |  |  |  |  |  |  |  | m* |  | m* |
| A4 |  | m | m |  | f* |  |  |  |  |  |  |  |
| XY | m* |  |  |  |  | m |  |  | f |  |  |  |

Interaction effects in best model:

A1:n genes:n RNA TEs (- ns), female-biased flower expression:female distortion (+ ns)

A2: n genes: flower male biased genes (+), male distortion: flower male biased genes (+ ns)

A3: position window:n genes (+ ns), position window:female distortion (+ ns)

A4: position window:n DNA TEs (-), leaf-expressed genes:n DNA TEs (+ ns)

XY: n genes:flower-expressed genes (-), n genes:pollen-biased genes (+)

#### Table S7

Full model results, including all coefficients, test statistics, and formulae, for all models. (Attached.)

### Genome distribution of expression

##### Figure S1

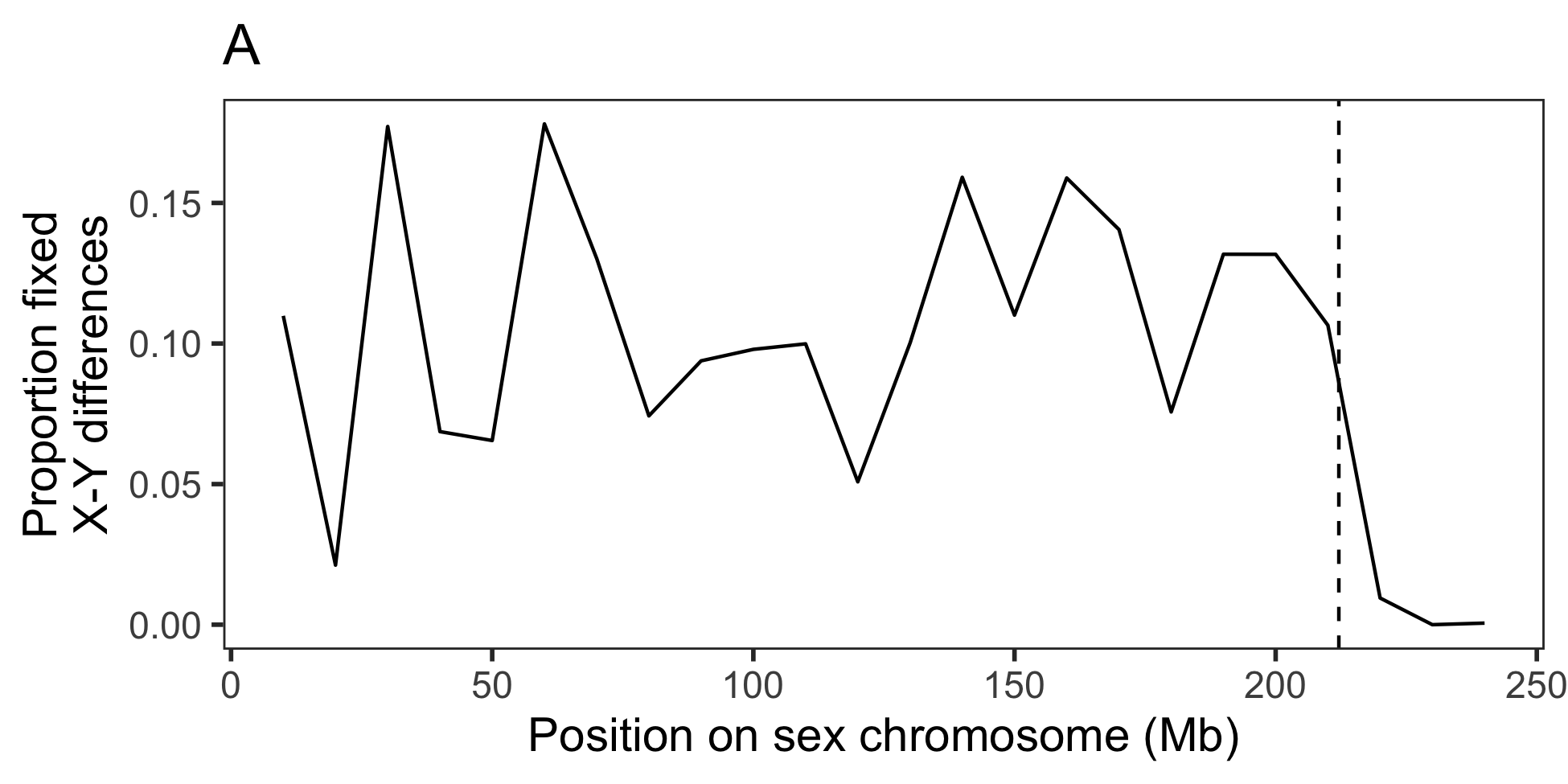

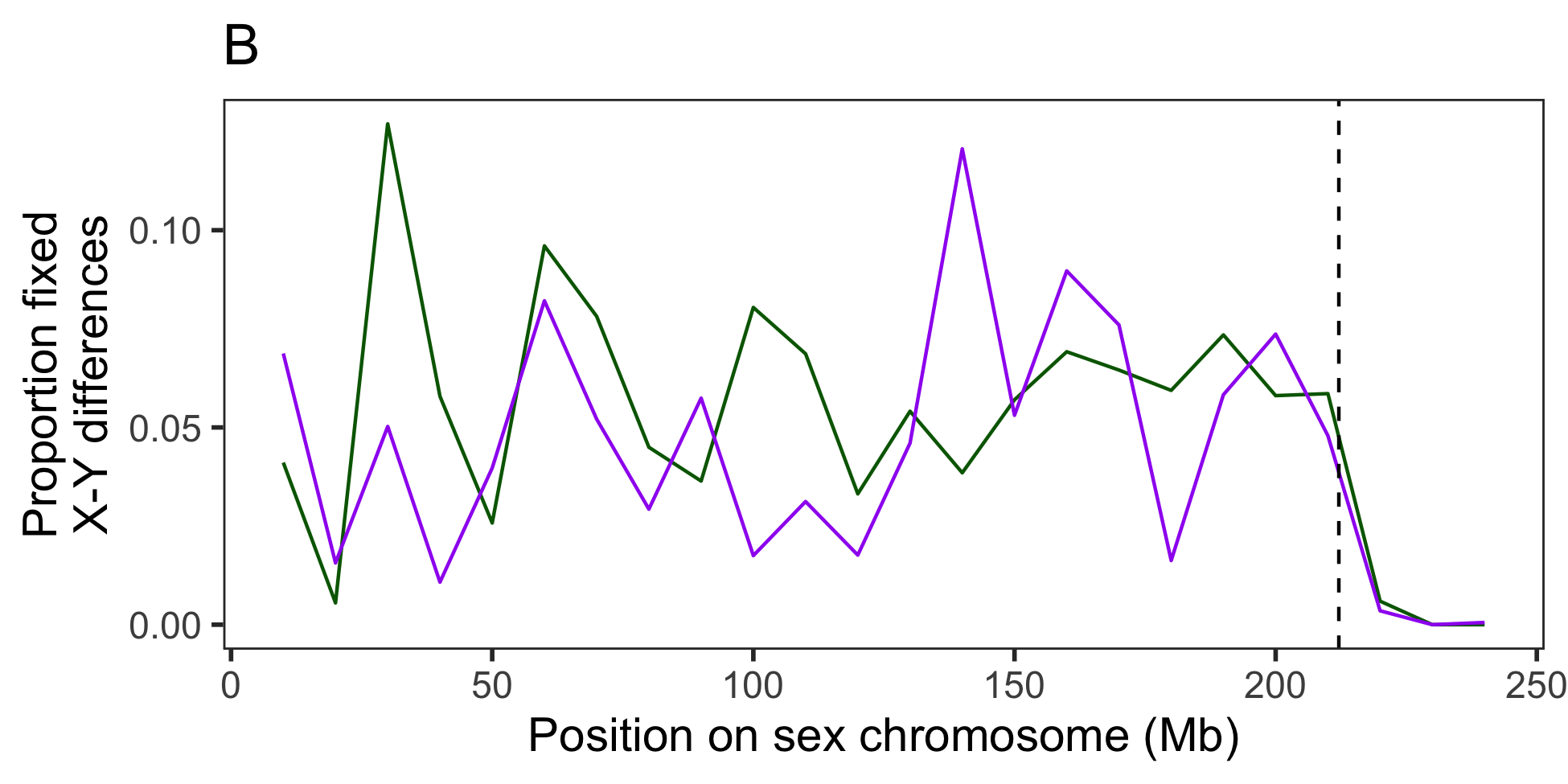

Figure S1- Proportion of SNPs fixed between X and Y chromosomes based on polymorphism analysis from transcriptome data of the XY cytotype [[1]](https://paperpile.com/c/IcnSpP/XAf1). Dashed line indicates the start of recombination in males at 212.1Mb. A. Proportion of biallelic SNPs where all males are heterozygous and all females are homozygous. B. Results separated by whether the reference assembly base is the X sequence (green-all females homozygous reference), and the reference assembly base is the Y sequence (purple- all females homozygous non-reference). Note that since this assembly is from a male, the sex chromosome represents a chimeric assembly between the X and Y sequence.

##### Figure S2

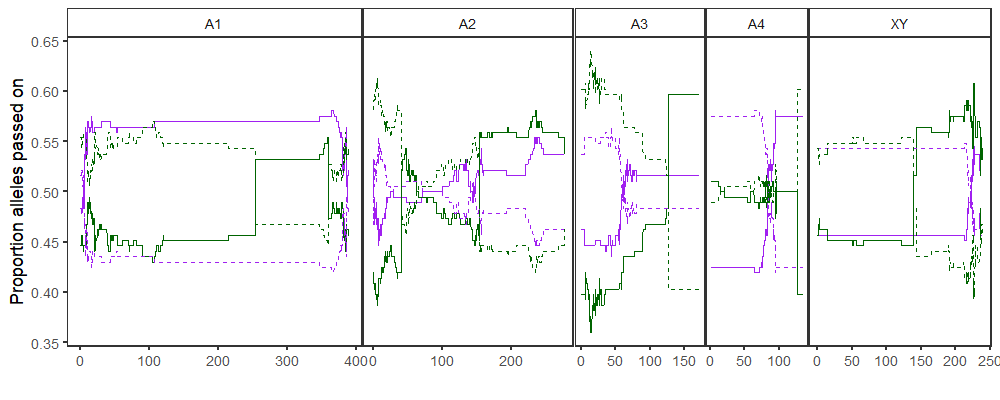

Figure S2 - inheritance proportions of paternal and maternal alleles across all chromosomes. Purple: alleles inherited from father. Green: alleles inherited from mother. Solid line: allele 1. Dashed line: allele 2.

##### Figure S3

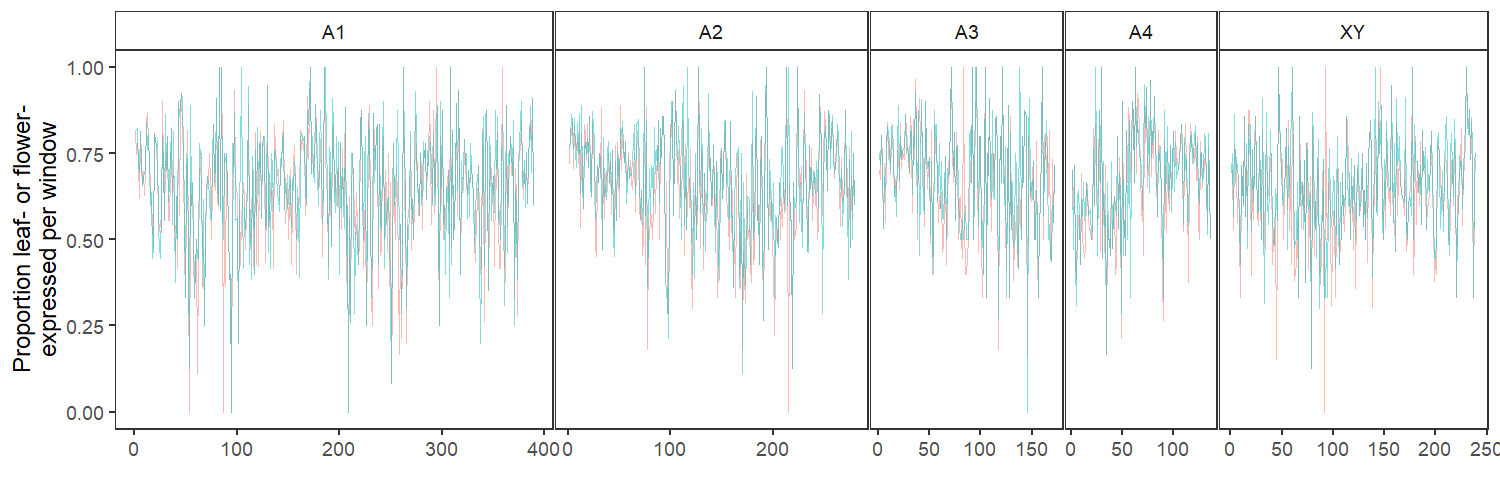
Figure S3: proportion of genes per window that are expressed in leaf (cyan) and flower (coral) tissue

##### Figure S4

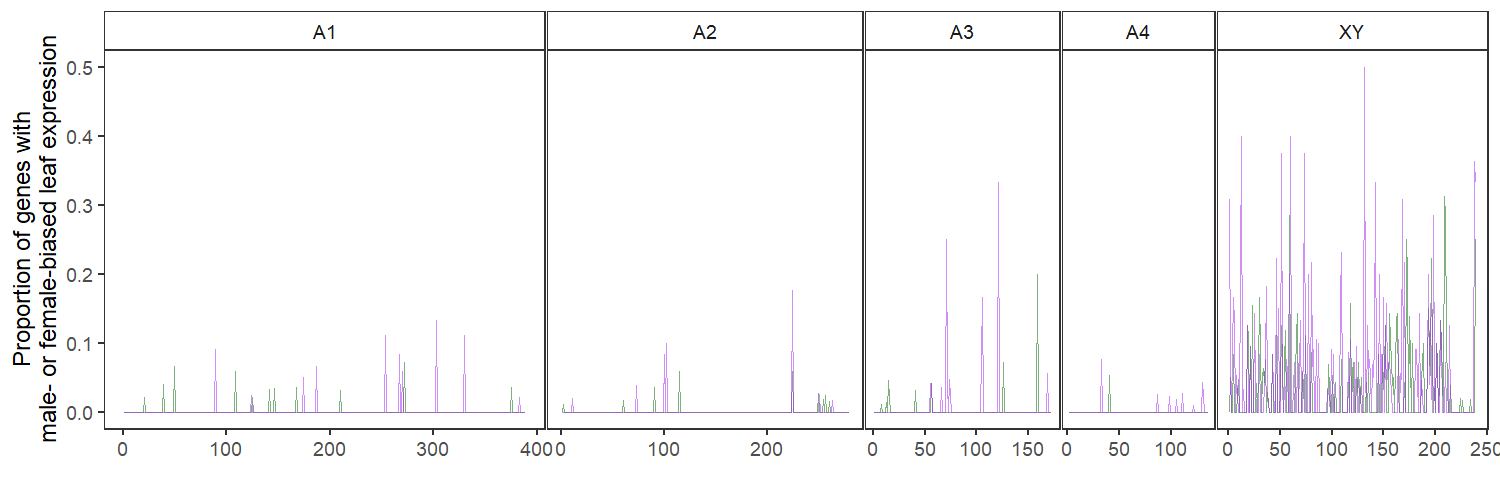

Figure S4: proportion of genes per window with male-biased (purple) and female-biased (green) expression in leaf tissue

##### Figure S5

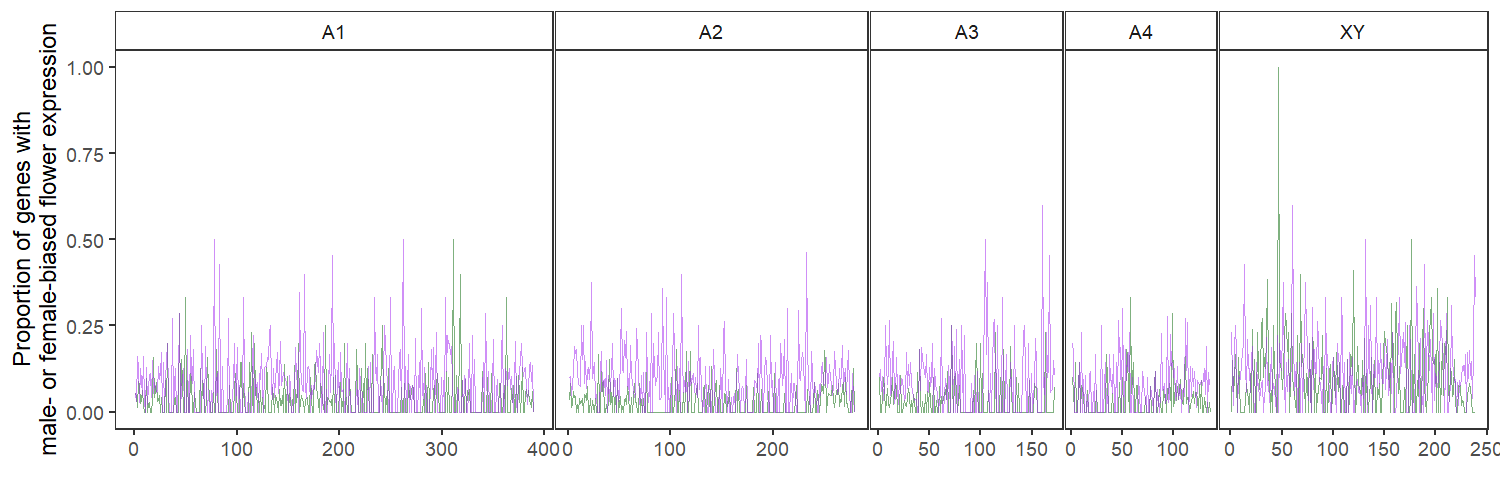

Figure S5: proportion of genes per window with male-biased (purple) and female-biased (green) expression in floral tissue

##### Figure S6

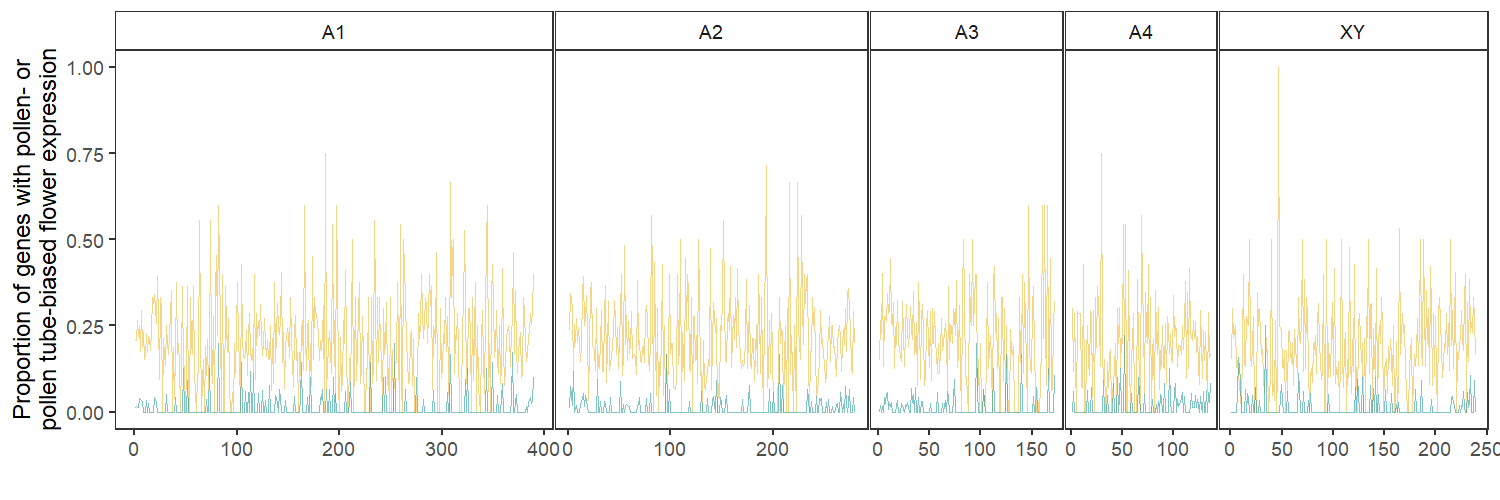
Figure S6: proportion of genes per window with pollen-biased (yellow) and pollen tube-biased (blue) expression.

### Correlations with crossover density

##### Figure S7

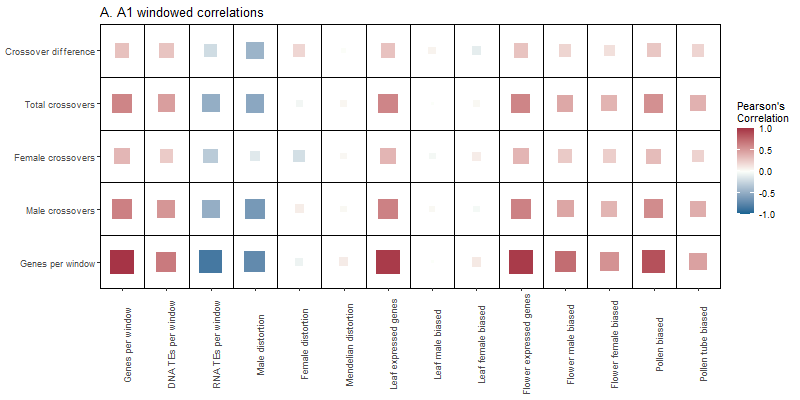

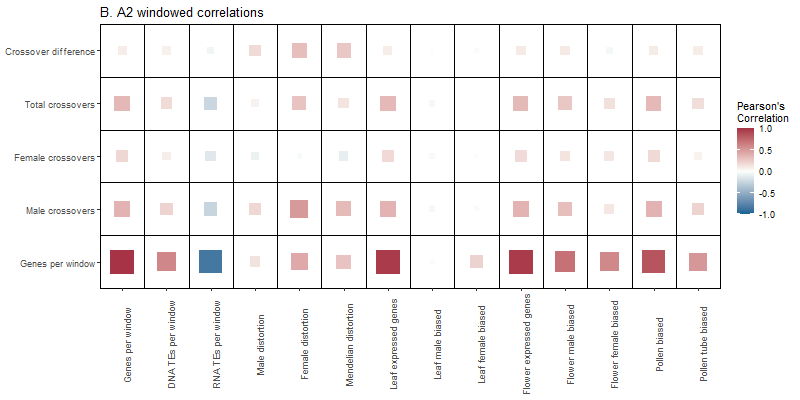

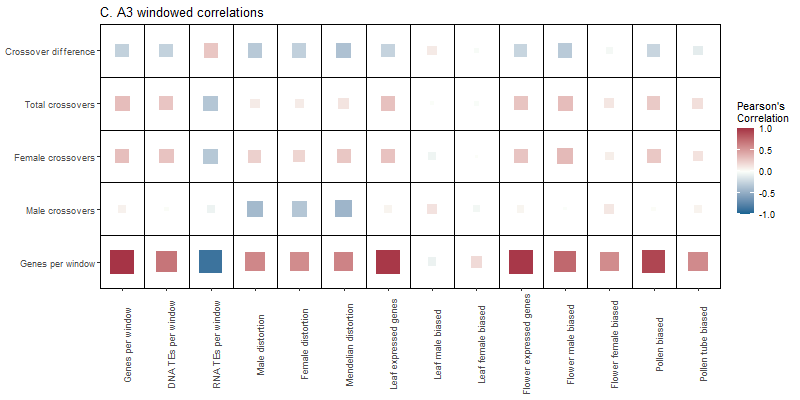

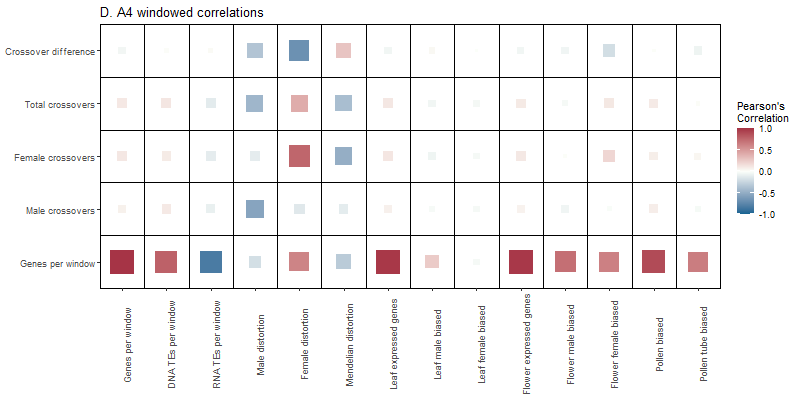

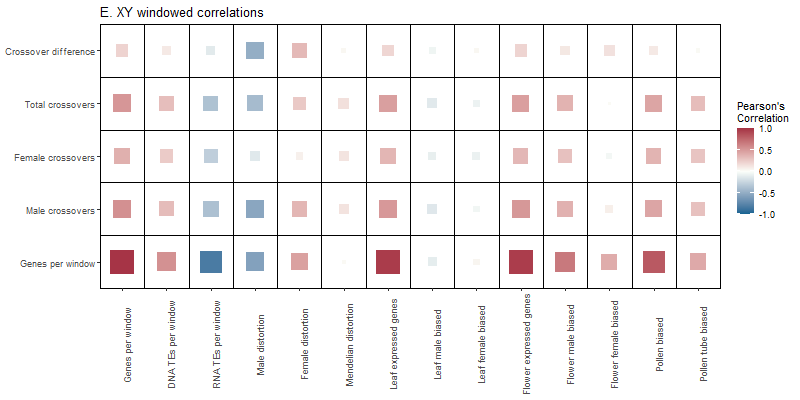

Figure S7: Correlations among genome window characteristics for each chromosome in *Rumex hastatulus*. Colours and sizes correspond to the strength and direction of the correlations. A: A1. B: A2. C: A3. D: A4. E: XY.

##### Figure S8

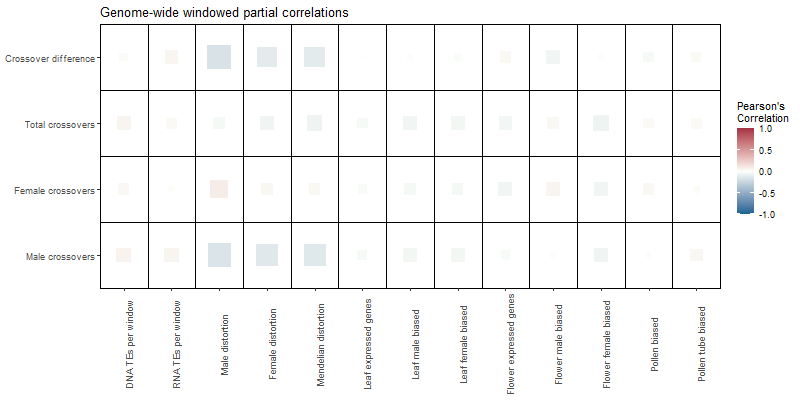

Figure S8: Partial correlations controlling for gene density among genome window characteristics across the whole genome in *Rumex hastatulus*. Colours and sizes correspond to the strength and direction of the correlations.

##### Figure S9

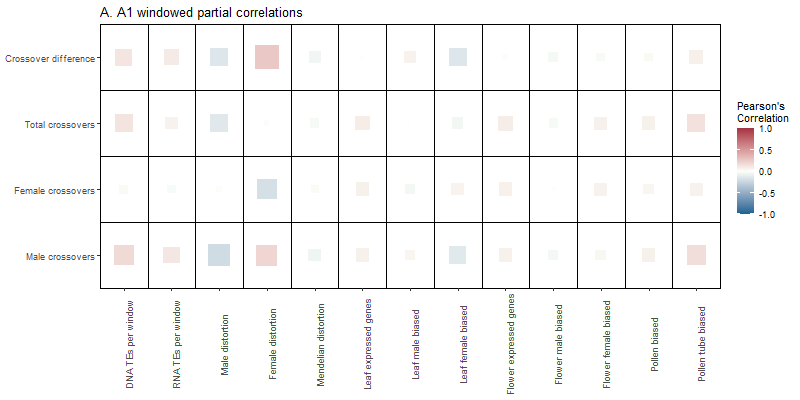

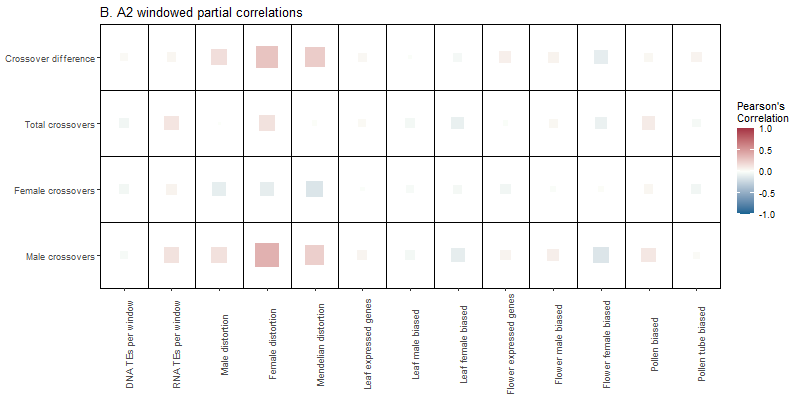

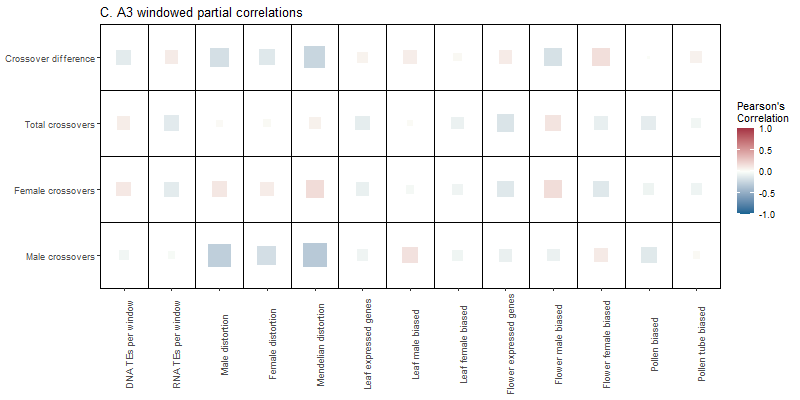

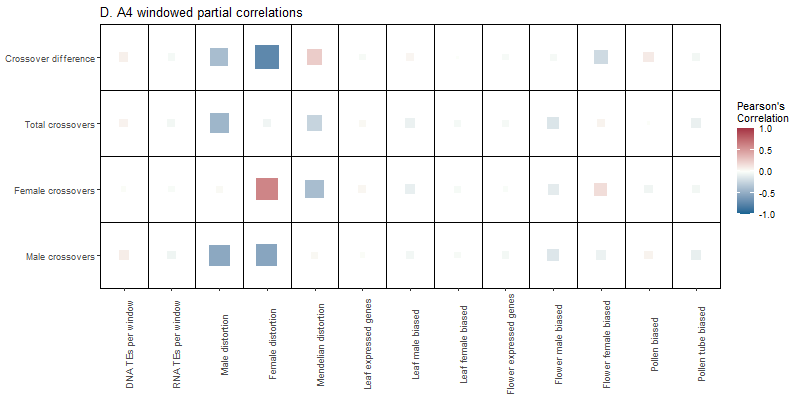

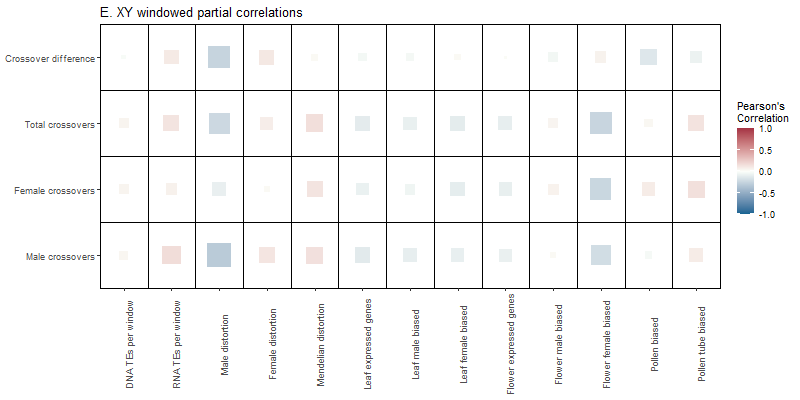

Figure S9: Partial correlations controlling for gene density for each chromosome in *Rumex hastatulus*. Colours and sizes correspond to the strength and direction of the correlations. A: A1. B: A2. C: A3. D: A4. E: XY.

1.	Hough J, Hollister JD, Wang W, Barrett SCH, Wright SI. 2014 Genetic degeneration of old and young Y chromosomes in the flowering plant Rumex hastatulus. *Proceedings of the National Academy of Sciences* **111**, 7713–7718.
